# Type VI secretion phenotypic heterogeneity ensures trade-off between antibacterial activity and resistance in Enteroaggregative *E. coli*

**DOI:** 10.1101/2025.02.11.637775

**Authors:** Boris Taillefer, Florian Schattenberg, Thierry Doan, Susann Müller, Eric Cascales

## Abstract

The type VI secretion system (T6SS) is a nanoweapon deployed by bacteria to deliver effectors into target cells, allowing niche colonization or host invasion. In entero-aggregative *Escherichia coli* (EAEC), the expression of the T6SS gene cluster is responsive to iron starvation by the Fur regulator, and to Dam-dependent methylation. However, fluorescence microscopy recordings showed that the T6SS displays heterogeneous expression in a clonal population. The population is composed of three types of cells in a stable equilibrium: non producers (OFF), producers, and assemblers (ON). Single-cell analyses further showed that the activation state is transmitted to the progeny, but the overall distribution respects the equilibrium suggesting a genetic control of heterogeneity. Mutagenesis of Fur-binding boxes and GATC sites revealed that specific promoter elements control this heterogeneity. Phenotypic characterization of the ON and OFF subpopulations demonstrated that ON cells eliminate susceptible target cells, whereas OFF cells survive attacks from T6SS-defensive bacteria. The heterogeneity of EAEC T6SS is therefore an attenuation strategy conferring a trade-off between antibacterial efficiency and survival in polymicrobial environments.

## INTRODUCTION

The type VI secretion system T6SS is a nanoweapon using a spring-like contractile mechanism to deliver effectors into target cells (1). By injecting toxins into bacterial rivals and/or eukaryotic cells, the T6SS is a key determinant in microbial communities and in pathogenesis (2). The T6SS is widespread in Gram-negative bacteria including bacteria living in various environments such as respiratory tract opportunistic pathogens, intestinal pathogens and commensals, plant-associated symbionts and pathogens, and soil or seawater bacterial species (3,4). T6SS gene clusters are therefore finely tuned by a broad variety of regulatory mechanisms. T6SSs are often responsive to physical, chemical and nutritional parameters such as cellular density, pH, temperature, salinity, viscosity, nutrient availability, host environmental cues and stresses via quorum sensing, two-component systems, alternative sigma factors or global regulators (5,6).

In EAEC, the expression of the Sci-1 T6SS gene cluster is controlled by iron levels through the ferric uptake regulator Fur, and by Dam methylation (7). The promoter P*sci1* is comprised of two Fur boxes (proximal F1 and distal F2 boxes) and three GATC (G1, G2 and G3) sites, with F1 and G1 overlapping the −10 of the transcription. This architecture creates a competition between Fur, the Dam methyltransferase and the RNA polymerase (RNAP) for the promoter, which will depend on iron availability. In iron-rich conditions, Fur-holo binds the promoter, prevents G1 methylation by Dam and consequently represses the expression of the T6SS gene cluster (OFF state). In iron-starved conditions, Fur-apo loses affinity for the promoter and allows Dam to methylate GATC sites excluding Fur from the promoter, leading to the induction of the T6SS expression (ON state) (7).

In several cases, such a promoter architecture involving competition between a transcriptional regulator and a DNA methyltransferase has been shown to produce epigenetic regulations responsible for phenotypic heterogeneity (8). Interestingly, while never described for a T6SS, fluorescence microscopy recordings using translational fusions to T6SS core components suggested heterogeneous expression in enteroaggregative *Escherichia coli* (EAEC) (9–15). Phenotypic heterogeneity is defined as the presence of different traits in a clonal population (i.e., genetically identical cells in a population). Phenotypic heterogeneity is produced by the differential gene expression from cell to cell, thus resulting in two, or more, subpopulations with a different expression state for a phenotypic trait (8,16–18). Phenotypic heterogeneity can arise from genetic (switch, rearrangement) or non-genetic (DNA methylation, feedback-loop, stochastic molecule partitioning) variations and serves two main adaptive strategies: bet-hedging and division of labor (19). The bet-hedging strategy allows one of the subpopulations to be pre-adapted to potential environmental changes, thus increasing the survival probability. It also permits to express a beneficial trait by optimizing the population fitness if the trait has a disadvantageous cost at the same time or in other conditions. An example of a bet-hedging strategy is the antimicrobial peptide resistance exhibited by a small fraction of the population in *Photorhabdus laumondii*, which is necessary for the invasion success of the larvae host (20). In the division of labour strategy, the different subpopulations accomplish specific tasks independently and hence increase the system functionality (21). A striking example of a division on labour strategy is found in the social bacterium *Myxococcus xanthus*, in which the heterogeneous expression of several genes enables the population to organize and form fruiting bodies that are essential for spore dispersal and population survival during nutrient starvation (22). Another well-characterized example is the *Salmonella enterica* Typhimurium type III secretion system (T3SS), which heterogeneous expression is producing ON cells capable of invading host cells and OFF cells dividing more rapidly to enable successful infection (23). In this case, the ON/OFF subpopulation equilibrium is critical for pathogenicity since a slight change of the ratio reduces the infection success (24). While more and more studies are describing phenotypic heterogeneity in several bacterial trait, the role and the ecological impact of such a phenomenon remains poorly understood (25).

The observed phenotypic heterogeneity of the EAEC T6SS questions the fitness cost of such an apparatus, which was speculated as energetically expensive for the cell (4,26). Such a fitness cost would explain the complex regulatory mechanisms that usually control T6SS expression and support the hypothesis that heterogeneous expression may solve growth defects due to T6SS expression and action (27). However, it was recently shown that T6SS production and assembly is not energetically expensive in EAEC, *Vibrio cholerae* and *Vibrio fischeri* (28–30). Moreover, these studies focused on the reproductive success in simple *in vitro* conditions and could miss specific and more complex *in vivo* conditions in which T6SS production and/or activity may impair the population fitness. Indeed, a fitness defect for T6SS^+^ *Campylobacter jujeni* cells was observed during bile salt exposition when competing against *E. coli* (31). Similarly, experimental evidence suggest a fitness cost for *Bacteroides fragilis* expressing the T6SS in the complex mice gut environment but not *in vitro* (32).

In this work, we characterize the origin and the role of phenotypic heterogeneity of the T6SS *sci1* gene cluster in EAEC 17-2. Using fluorescent reporters, confocal microscopy and microfluidic, we showed that OFF and ON subpopulations are at equilibrium in iron starved conditions. Using cell sorting to isolate ON or OFF subpopulations, we observed that homogeneous subpopulations return to the heterogeneous equilibrium in about ten generations. We then deciphered the molecular basis of this heterogeneity by using mutagenesis of regulator binding sites, demonstrating that binding of Fur to the distal F2 Fur box and methylation of the G3 GATC site are key determinants of P*sci1* heterogeneity. We finally used a competition assay to address the role of heterogeneity. Our results suggest that EAEC T6SS heterogeneity is an attenuation strategy in a polymicrobial environment. While the role of ON cells is to eliminate rivals, we found that OFF cells increase survival by reducing tit-for-tat responses by defensive T6SS^+^ bacteria.

## RESULTS

### *sci1* expression is heterogeneous in a clonal population of EAEC

The EAEC Sci1 T6SS is controlled by iron levels through Fur-mediated repression. T6SS expression is therefore repressed in iron rich conditions, such as in LB medium, and induced in iron depleted conditions, such as in M9/glycerol minimal medium (*sci1*-inducing medium, SIM, 7). Using the T6SS translational reporter strain *tssB-sfGFP* (B-GFP) as a model to study Sci1 T6SS sheath dynamics in enteroaggregative *E. coli* (EAEC), we observed heterogeneity in a clonal population by fluorescence microscopy in SIM medium (**Figure 1A, left panel**). Three subpopulations could be observed: cells that do not produce TssB-sfGFP (no fluorescence), cells that produce TssB-sfGFP (diffuse fluorescence) and cells that produce TssB-sfGFP and assemble sheaths. These results suggest that T6SS production is subjected to two heterogeneous regulations, at the transcriptional or translational level (production vs no production) or at the post-translational level (assembly vs no assembly). To gain further information on the transcriptional/translational heterogeneity, we engineered a transcriptional reporter strain in which the sequence encoding the sfGFP was inserted between the *tssC* and *tssK* genes (C-GFP-K). This reporter showed that the EAEC Sci1 T6SS is heterogeneously expressed (**Figure 1A, right panel**), with a bimodal distribution of the fluorescence in the population, defining two major subpopulations, OFF (*∼*40%) and ON (*∼*60%), with low and high fluorescence, respectively (**Figure 1B**, **Figure S1**). It is noteworthy that the two peaks significantly overlap, suggesting that intermediate and possibly transient activation states exist **(Figure 1B**). The existence of OFF and ON subpopulations, and of intermediate states was further confirmed by cell sorting (**Figure S4A**, see below).

**Figure 1.**
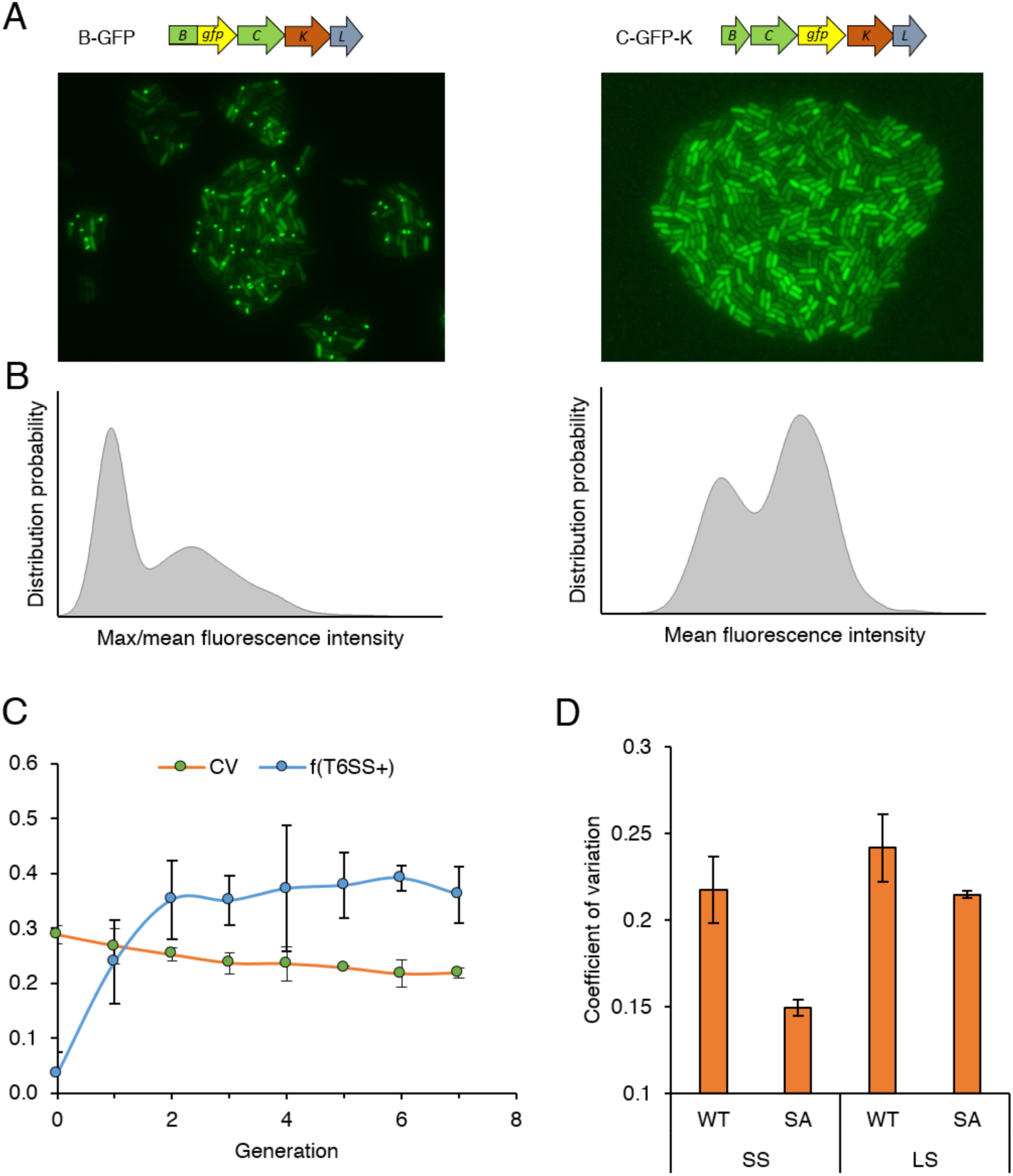
The phenotypic heterogeneity of the T6SS expression and assembly reaches an equilibrium which is stable over generations. **A**. Representative fields of B-GFP and C-GFP-K heterogeneous expression observed under the confocal microscope. Images shown are the fluorescent channel only. **B**. Representative graph of heterogeneous T6SS activity and expression distribution in a clonal population of B-GFP (left) and C-GFP-K (right), respectively. B-GFP fluorescence is expressed in max/mean fluorescence intensity ratio to distinguish cells with bright T6SS apparatus. Low max/mean fluorescence indicates no sheath whereas ratio > 2 indicate the presence of at least one sheath. Y-axis represents the distribution probability of the fluorescence intensity measured in each cell of the population. **C**. Coefficient of variation (CV) of C-GFP-K population and T6SS^+^ cell frequency (f(T6SS^+^)) of B-GFP population (y-axis) in function of the generation number (x-axis). T6SS^+^ cell frequency was calculated as the number of cell with a max/mean fluorescence intensity > 2. Approximatively 2.000 cells were analysed for each generation from at least 3 independent replicates. **D.** Coefficient of variation (y-axis) of the ancestral population (C-GFP-K strain, WT) compared to the SIM-adapted population (SA). Prior to the measurement, WT or SA populations were successively cultured (preculture and subculture) in SIM (SS) or in LB and then in SIM (LS). More than 2,500 cells were analysed for each conditions, from 3 independent replicates).

To determine the level of heterogeneity, we monitored the coefficient of variation (CV), i.e., the ratio between the standard deviation of the fluorescence represented by values from each cell with the mean fluorescence of the population (CV=SD/µ). The higher is the CV, the higher is the heterogeneity; the lower is the CV, the higher is the homogeneity. We observed that the ON/OFF equilibrium is stable over generations, as shown by the stability of the coefficient of variation or the T6SS^+^ frequency in the population (**Figure 1C**, **Figure S2**). Only a long-term culture (approx. 50 generations) in SIM achieved by successive passages led to a shift of the equilibrium to a more homogeneous population (SS condition, **Figure 1D**). This homogeneity, however, was lost after a round in repressive medium to come back to a WT value (LS condition, **Figure 1D**).

### Heterogeneous expression is stable and transmissible

To better understand how this heterogeneity emerges and spreads in the population, we performed single-cell analyses. C-GFP-K cells were grown until the equilibrium, diluted and plated on a microfluidic chip so that a single cell can be observed per field. Time-lapse experiments showed that a single cell can produce homogeneous OFF, homogeneous ON and heterogenous microcolonies (**Figure 2A**, **Figure S3**). Further analyses showed that heterogeneous and homogeneous ON microcolonies appears more frequently compared to OFF microcolonies, yielding an overall distribution such as what is observed in liquid culture (CV > 0.20, **Figure 2B**). We however observed a lower T6SS^+^ cell frequency during single-cell experiments which we could explained by a reduced growth due to the oxygen availability compared to liquid cultures (**Figure 2B**). The observation of homogeneous microcolonies suggests a transmission of the activation state from mother to progeny which frequency seems to be different for ON and OFF cells.

**Figure 2.**
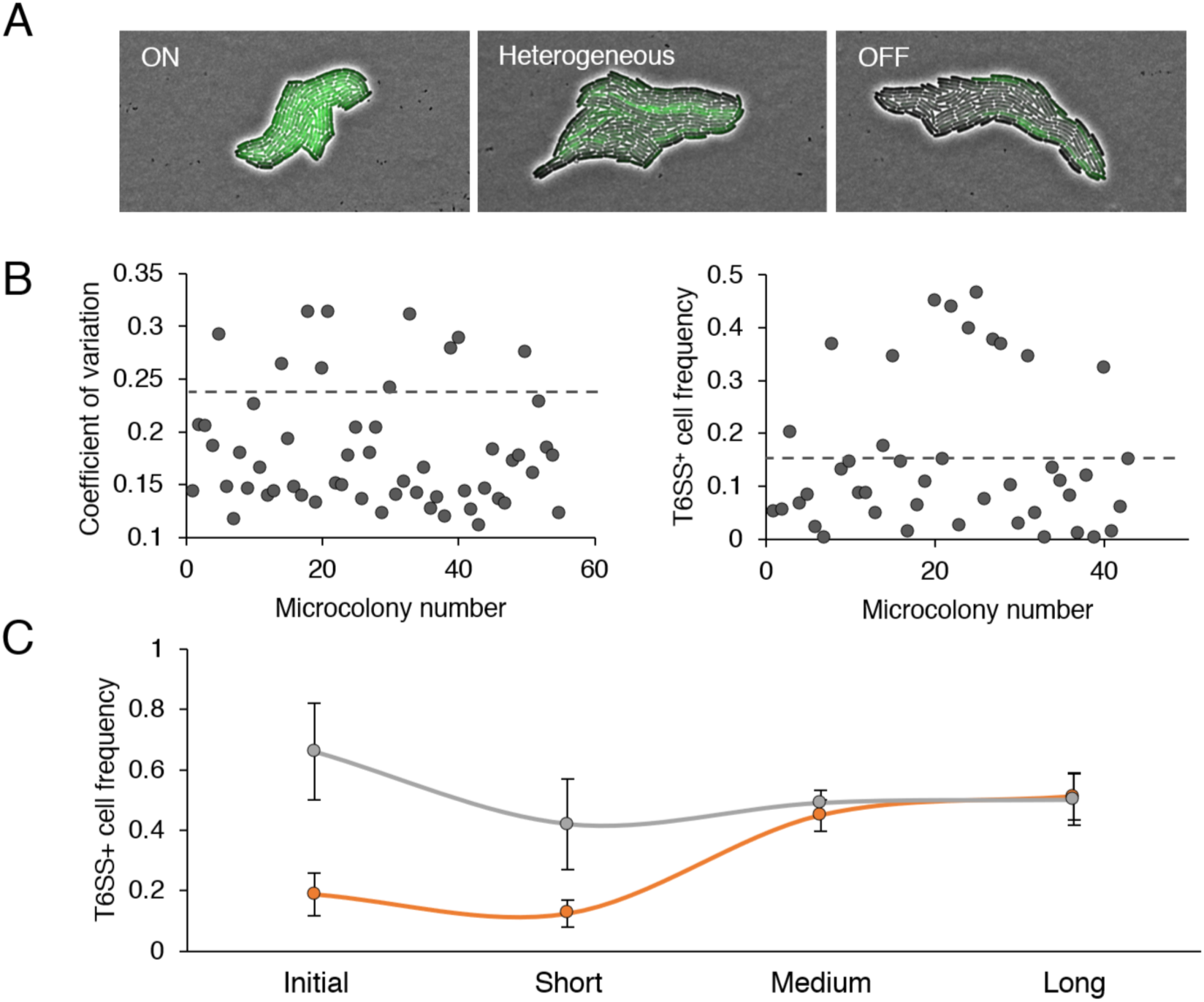
Single-cell experiments gave rise to 3 types of microcolony, but the overall distribution is heterogeneous. **A**. Illustration of the 3 types of C-GFP-K microcolonies (homogeneous ON, heterogeneous, homogeneous OFF). **B**. Distribution of the coefficient of variation (CV) in C-GFP-K (left) and of the T6SS^+^ cell frequency in B-GFP (right) in microcolonies from single-cell experiments. Average frequency and coefficient of variation is noted by the dotted line. C-GFP-K: 8,500 cells analysed (55 microcolonies from 4 independent replicate)s; B-GFP: .900 cells analysed (43 microcolonies from 3 independent replicates). **C**. Dynamic of T6SS^+^ cell frequency in sorted subpopulations at different timescale. Initial: frequency after sorting; Short: frequency in microcolonies (5 generations); Medium: primary liquid culture (15 generations); Long: secondary liquid culture (20 generations). Each experiment was performed at least 3 times.

To further characterize these subpopulations, ON and OFF cells were sorted by flow cytometry (**Figure S4A and S4B**). The ratio of OFF and ON cells was comparable to that observed by fluorescence microscopy. Single-cell experiments confirmed a transmission of the activation state in short-term with a majority of OFF and ON microcolonies formed by OFF an ON sorted cells, respectively (**Figure 2C, Figure S4C**). However, the OFF and ON subpopulations returned to the equilibrium in 10-15 generations when cultured in SIM medium (**Figure 2C**).

### Fur boxes and GATC methylation sites are responsible for P*sci1* heterogeneous expression

The recovery and the stability of the equilibrium suggests that T6SS heterogeneity is genetically controlled. The *sci1* promoter (P*sci1*) is comprised of multiple regulatory elements including two Fur boxes (proximal F1 and distal F2) and three GATC sites, which are target of the Dam methyltransferase (proximal G1 to distal G3) (**Figure 3A**). G1 is included within the F1 box, which overlaps the −10 element. G2 is located between the −10 and −35 elements, while F2 and G3 are located upstream the −35 (**Figure 3A**). Fur binding onto F1 and F2 was shown to repress *sci1* expression and to prevent G1 methylation. In contrast, G1 methylation was shown to reduce Fur binding, thus allowing transcription (7).

**Figure 3.**
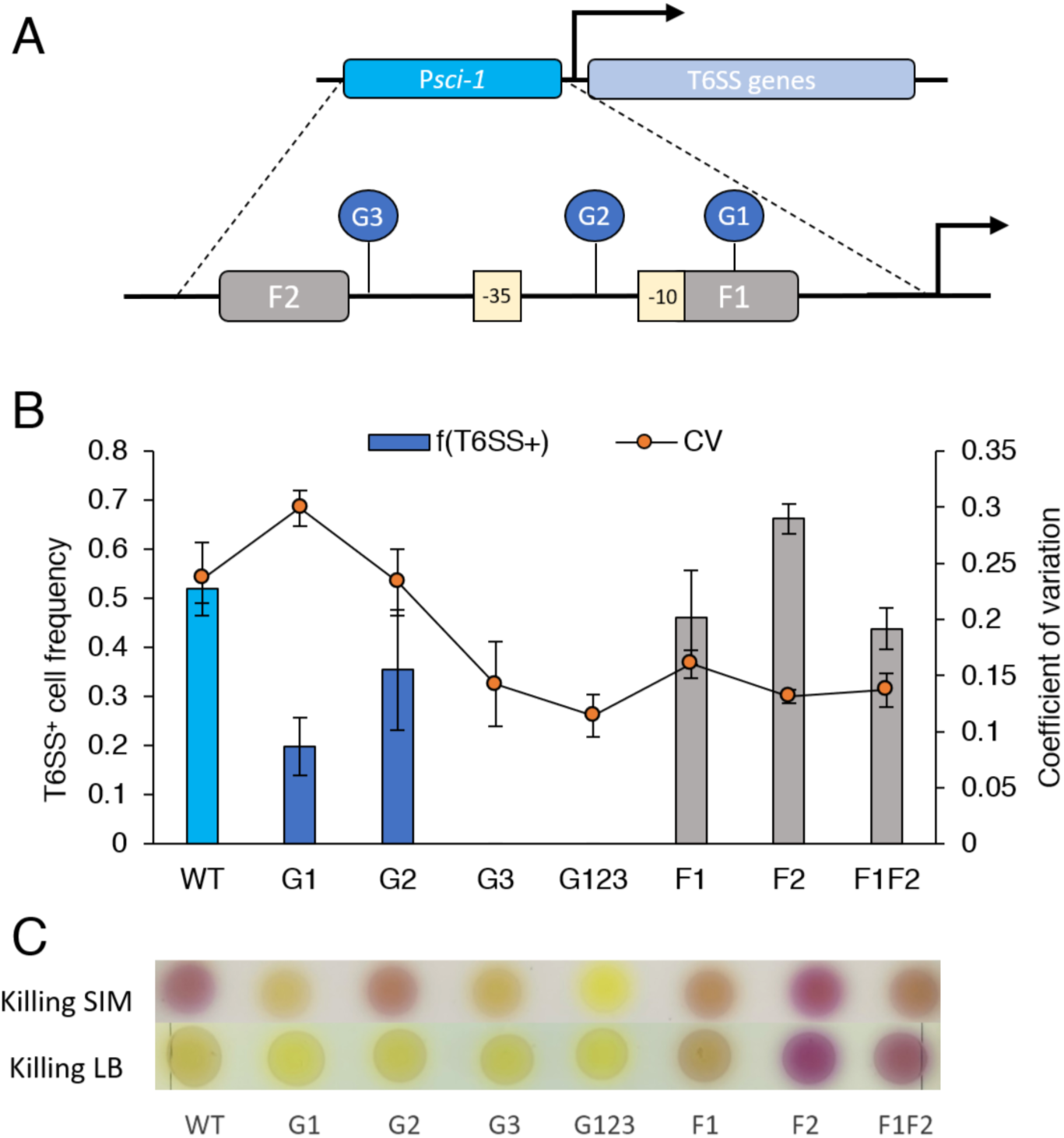
F2 and G3 are responsible for the heterogeneous expression. **A**. Schematic representation of the Sci1 T6SS promoter (P*sci1*). The promoter controls the *sci1* operon containing all T6SS essential genes. P*sci1* is composed of two Fur binding boxes (F1 and F2) and three GATC sites, which are the targets of the Dam methyltransferase (G1, G2 and G3). The F1 box includes the G1 site and overlaps with the −10 of transcription (7). **B**. T6SS^+^ cell frequency of B-GFP (bar plots, f(T6SS^+^)) and coefficient of variation(CV) of C-GFP-K (line points) EAEC cells and promoter variants. Data are the mean of 3 independent replicates. **C**. Competition assay of EAEC and promoter variants (attacker) against sensitive *E. coli* W3110 (recipient) in SIM (top) or LB (bottom) medium. The LAGA method is used where colour intensity reflects the attacker killing activity (33), from yellow (no killing) to purple (intense killing).

To define the impact of these regulatory components on Sci1 heterogeneity, we engineered chromosomal mutations to disrupt the different elements. Mutagenesis of GATC sites to prevent adenine methylation confirmed that these sites are involved in P*sci1* activation as G1, G2, G3 and the triple GATC mutant G123 presented decreased sheath numbers (**Figure 3B**). Mutagenesis of the Fur boxes also confirmed previous studies on their repressive role as F1, F2 and F1F2 mutations alleviated repression and increased sheath numbers. When introduced into the C-GFP-K strain, Fur box mutants presented a homogeneous ON distribution, while G3 and G123 mutants had a homogeneous OFF distribution (**Figure 3B**). Although G1 and G2 mutants were heterogeneous, the ON/OFF equilibrium was different from the WT (**Figure 3B, Figure S5, Figure S6**). Taken together, all the mutations impacted the promoter activity as well as the subpopulation equilibrium, suggesting that the promoter architecture favours a specific heterogeneous expression.

To determine the impact of these mutations on T6SS activity, we conducted antibacterial assays. **Figure 3C** shows that the T6SS activity of the different mutants against sensitive W3110 recipient cells correlates with the T6SS^+^ frequency: the G1 and G2 mutants presented an impaired T6SS activity, whereas G3 and G123 mutant cells did not show any antibacterial activity in SIM (**Figure 3C**). In contrast, the F2 and F1F2 mutations increased T6SS activity compared to the WT and F2 and F1F2 mutants eliminated target cells in repressive, iron-rich LB conditions (**Figure 3C**). These results therefore agree with the postulate that ON cells deploy T6SS to kill competitors whereas OFF cells do not express T6SS.

### T6SS heterogeneity is an attenuation strategy against defensive T6SS

As the F2 and G123 mutants simulate the ON (T6SS^+^) and OFF (T6SS^-^) subpopulations respectively, we conducted experiments to determine the role of this phenotypic heterogeneity, and notably of OFF cells. The T6SS has long been thought to be costly for the cell. A costly machine model agrees with a heterogeneous expression, as the population benefits from ON cells for the function of the machine and from OFF cells for counteracting the growth defect associated to the production, assembly or activity of the machine. However, we recently showed that no energetic fitness cost is entailed by the T6SS in EAEC (29). Using homogeneous strains engineered in this study, we still did not observe any growth impact in liquid culture for T6SS overexpression (F2, ON) or complete inhibition (G123, OFF) (**Figure S7A**). Taken together, these results suggest that the T6SS growth cost is negligeable in EAEC.

Another hypothesis that could explain T6SS phenotypic heterogeneity is the suicide strategy, where a subpopulation sacrifices itself (or is killed) in nutrient starved conditions to support the growth of the other cells. In the case of the T6SS, one could hypothesize that OFF cells serve as a nutritional reservoir for ON cells in single species culture. However, we never recorded kin killing in EAEC as exemplified by a competition assay that did not impact EAEC WT mortality when facing the ON homogeneous F2 strain (**Figure S7B**), likely because of the basal expression of the *tli1* gene encoding the immunity protein due to the P*_4532_* internal promoter (34).

Another hypothesis for T6SS heterogeneity is an attenuation into the host. During host infection, the immune system recognizes virulence factors to neutralize pathogens. An attenuation strategy, such as LPS modification in *Burkholderia pseudomallei* (35) or antigenic variation in *Borrelia burgdorferi* (36), allows to escape the immune system. This hypothesis was however infirmed in the wax moth model *Galleria melonella*. We observed that, while the homogeneous ON population had a higher infective success, OFF cells failed to successfully colonize *Galleria* (**Figure S7C**).

In the host gut, EAEC encounter competitors, including T6SS^+^ species. Some bacterial species have been shown to express and activate defensive T6SSs, which use a tit-for-tat retaliation mechanism to respond to T6SS attacks (37). A heterogeneous expression could be thus an attenuation strategy to diminish the defence intensity and to better survive. To test this hypothesis, we used the sensitive W3110 strain as recipient and confirmed that G123 OFF cells alone do not eliminate W3110. However, increasing the ON/OFF ratio resulted in a proportional increase in the antibacterial activity of the population, indicating that the T6SS activity strength is adjustable by controlling the fraction of cell expressing the system in the population (**Figure 4A**). In contrast, when the defensive T6SS^+^ *P. aeruginosa* PAO1*ΔretS* strain was used as recipient, we observed that increasing the ON/OFF ratio leads to increased EAEC mortality (**Figure 4B**). Taken together, these results suggest that ON cells are detrimental for the population survival in a polymicrobial environment, notably against a defensive T6SS^+^ bacterium.

**Figure 4.**
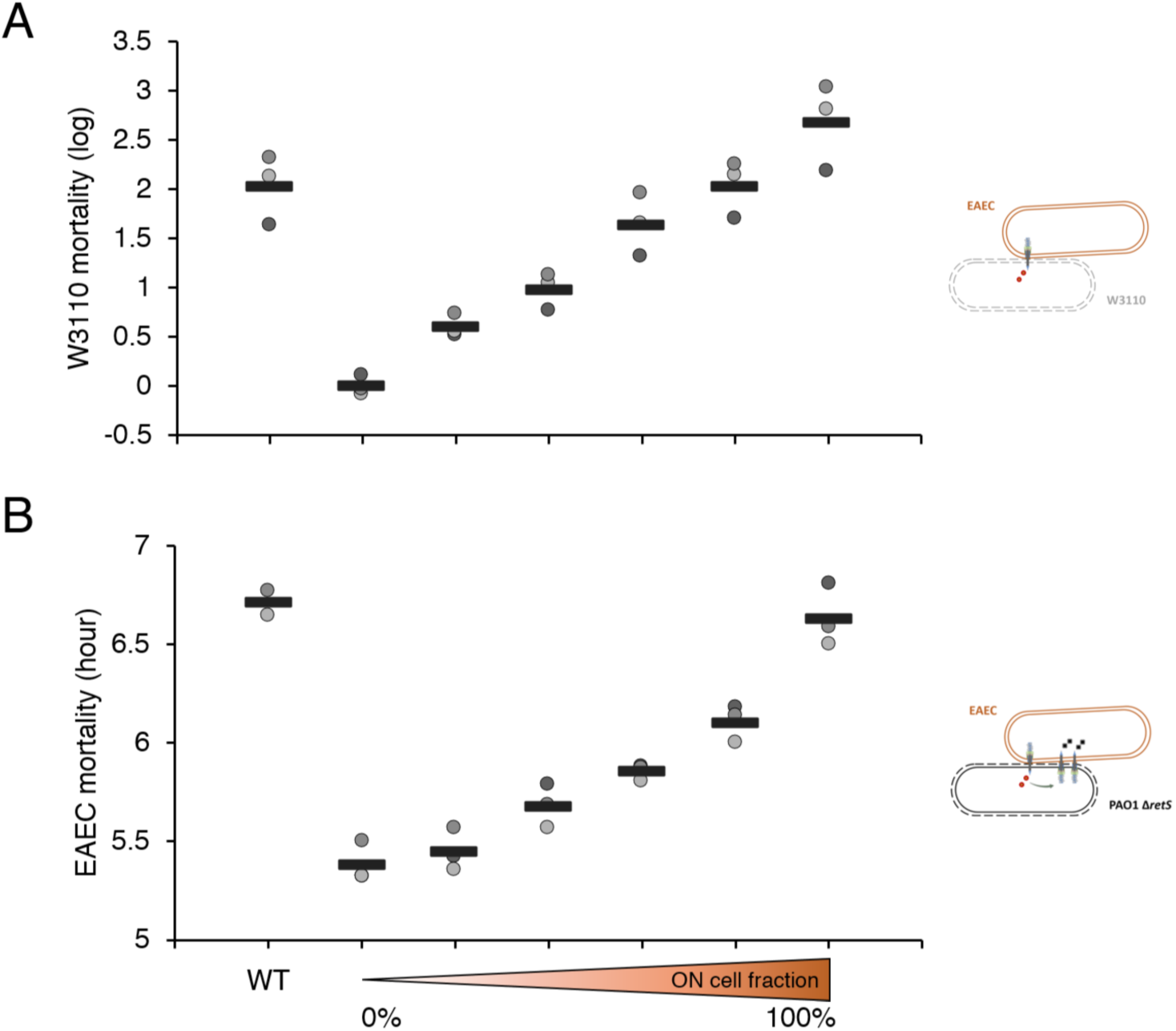
Heterogeneity controls the T6SS activity strength and answers a trade-off between killing efficiency and survival against T6SS^+^ defensive bacteria. **A**. Competition assay between wild-type (WT) EAEC or increasing T6SS ON/OFF ratio (from 0% to 100% ON fraction) and *E. coli* W3110 recipient. F2 and G123 mutants were used as ON and OFF cells, respectively, and combined in different ratios to compose different heterogeneity equilibriums. Recipient W3110 mortality was measured by the survivor’s growth recovery method (33) and log differences were determined thanks to a standard curve. Data represents 3 independent replicates. **B**. Competition assay between WT EAEC or increasing T6SS ON/OFF EAEC ratio and the T6SS^+^-defensive bacterium *Pseudomonas aeruginosa* PAO1*ΔretS*. EAEC mortality was measured by the survivor’s growth recovery method without standard curve but using the time for the population to recover. Data represents 3 independent replicates.

## DISCUSSION

In this work, we described and characterized for the very first time a phenomenon of T6SS phenotypic heterogeneity. We showed that the EAEC Sci1 T6SS is heterogeneously expressed when grown in iron-limited *sci1* inducing minimal medium. This heterogeneity is at a stable equilibrium with about 40% of OFF and 60% of ON cells. Although stable, this equilibrium is not locked, and ON and OFF states are reversible, indicative of an epigenetic control for T6SS heterogeneous expression (8). When grown in iron-rich LB medium, the population is homogeneously OFF, but dilution into minimal medium allows to reach the equilibrium. Similarly, culturing sorted ON and OFF cells regenerated the equilibrium in about 10-15 generations.

We then defined the genetic determinants responsible for heterogeneity. We demonstrated that the P*sci1* regulatory elements are also responsible for heterogeneous emergence, indicative of a genetic control. Mutagenesis of the Fur boxes induced T6SS expression, which is constitutively expressed, with homogeneous ON cells. In contrast, mutagenesis of the G3 GATC sites yields a homogenous OFF population, in agreement with previous results demonstrating competition between Fur and Dam at the P*sci1* promoter (7). Therefore, Fur regulates the expression of the Sci1 T6SS gene cluster, while the methylation state of the GATC sites modulates Fur binding, allowing heterogeneous expression from cell to cell. Interestingly, individual mutation of each of the promoter elements produced different level of heterogeneity and T6SS^+^ cell frequency. This indicate that a heterogeneous equilibrium could be attain by a specific promoter sequence, depending on the presence of Fur and Dam sites, as well as the affinity of the Fur repressor for its sites (38). These promoter architectures could be prone to natural selection to set a specific and adaptive T6SS ON/OFF ratio. This hypothesis is consistent with a recent study modelling the expression profile of several promoter sequence variations, indicative of the impact of small changes on regulator affinity and processivity (39).

We then show that, as expected, ON cells producing an active T6SS have antibacterial activity. OFF cells do not solve a potential T6SS cost but instead limits attacks by responsive T6SS^+^ bacteria. EAEC T6SS heterogeneity is therefore a trade-off strategy that solves a conflict between two contradictory behaviours: the ON state is critical for eliminating sensitive recipients but also a source of deadly responses from armed competitors. Indeed, in polymicrobial communities such as the gut microbiota where hundreds of species cohabit, this strategy may allow to efficiently kill competitors, but also to protect from counterattacks.

Defensive behaviours against T6SS attacks begin to be uncovered. Several mechanisms have been evidenced in the recent years, such as capsule production, protective aggregation, deployment of a defensive T6SS, membrane and permeability modification, accumulation of immunity genes or toxin-target modification (40–45). Interestingly, each of these resistance mechanisms is employed at a high fitness cost for the cell, notably EPS production and membrane modification (46). We described here that the T6SS heterogeneous expression in EAEC attenuates its activity and thus decreases the defensive response of competitors. As such, we propose that T6SS heterogeneity could be considered as an indirect adaptive defensive mechanism that optimizes the cost for resistance.

In the model shown in **Figure 5**, we propose that a homogeneous OFF population is unable to invade the gut because of its inability to destroy competitors. A homogeneous ON population would also be excluded, because of the high defensive reciprocation from the resident microbiota. In contrast, the heterogeneous population colonizes successfully by eliminating competitors and by reducing the defensive response. In addition, due to the reversibility of the ON and OFF states, we propose that the OFF subpopulation serves as a source of ON cells to replace those eliminated by tit-for-tat retaliation.

**Figure 5.**
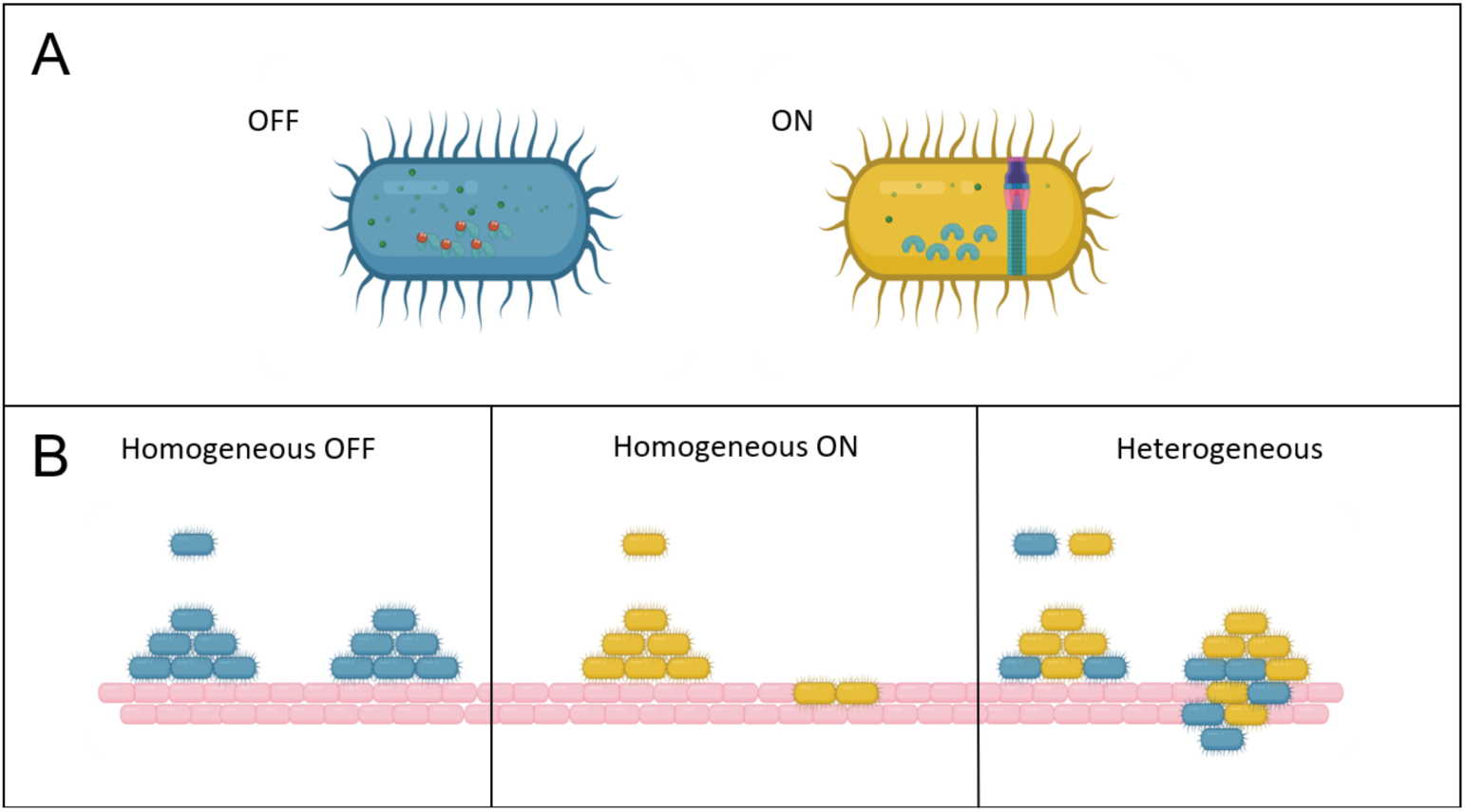
Proposed model for the T6SS phenotypic heterogeneity functional role. **A**. OFF and ON cells could differ from the intracellular iron content and thus the Fur activity. **B**. During the colonization process in the gut, the T6SS is useful to invade a niche. The OFF subpopulation is not able to invade the niche (left box). The homogeneous ON population can kill the bacteria encountered but sudden a high defensive response from resident bacteria (center box). A heterogeneous population attenuates the defensive response which optimizes the development of the population (right box). This figure was edited in Biorender.

Our results confirm that Fur is the main determinant of the regulation of the *sci1* gene cluster. Reversible Fur binding onto specific Fur boxes located in the promoter region controls the switch between the ON and OFF states (7). Mutation of the Fur boxes, preventing Fur binding, led to a constitutive ON state, even in presence of iron. T6SS heterogeneous expression therefore emanates by the probable modulation of Fur binding. Fur binds to Fur boxes once complexed to iron and is released when iron is limiting. Heterogeneity can therefore arise from heterogenous *fur* expression. This hypothesis is not supported by our observations that the transcriptional reporter fusion of P*fur* is homogenously expressed in EAEC (**Figure S8**). Instead, the competition between Fur and Dam suggests that stochastic methylation of the P*sci1* GATC sites modulates Fur binding and hence controls heterogeneity. The result of the competition between Fur and Dam will determine the activation state of the cell. Interestingly, the fluorescence microscopy and FACS analyses did not show two distinct peaks but rather a continuous distribution between OFF and ON cells, suggesting intermediates states. These intermediate states could be due to the dilution of the content of ON cells that switched to an OFF state, or to different expression levels due to hemi-methylated states. Despite the discovery of some genetic aspect explaining T6SS heterogenous emergence in EAEC (F2 and G3 elements), these elements are not sufficient to fully explain the heterogeneity dynamics. Additional criteria, such as Fur and Dam levels and dynamics, iron levels or methylation patterns could also play a role in T6SS heterogeneity. Microfluidic approaches and single-molecule tracking will be useful to better understand emergence of T6SS heterogeneity (47–49).

Fluorescence microscopy recordings of translation fusions to T6SS subunits showed that the T6SS is heterogenous in many species, including *Burkholderia thailandensis*, *Pseudomonas putida*, *P. fluorescens*, *Serratia marcescens*, *Salmonella enterica* Typhimurium*, Acinetobacter baylii* or *Vibrio fischeri* (50–54). The T6SS gene clusters in these species are under the control of very distinct regulatory mechanisms, such as xenogenic silencing, two-component systems or quorum sensing (6,55). It would be interesting to identify the molecular determinants underlying T6SS heterogeneity in these species that colonize different environments. Defining the function of heterogeneity would help to determine whether heterogenous expression answers different trade-off, division of labor or bet-hedging strategies. Since T6SSs are employed to kill bacteria in polymicrobial environment, we hypothesized that such an attenuation strategy by a heterogeneous expression might be conserved.

Finally, in addition to the heterogenous expression of the *sci1* gene cluster, imaging cells carrying the TssB-sfGFP translational fusion showed heterogenous assembly of T6SS sheaths. Further analyses are required to understand how T6SS assembly is activated in EAEC, and how assembly or activation is heterogeneously governed. One possibility is the control of stochiometric levels of T6SS subunits, which is crucial for proper assembly. It is worthy to note that the accurate expression/production of T6SS subunits is ensured by a second, internal promoter, P*_4532_*, that shares a similar architecture with P*sci1* (Fur and Dam sites overlapping the −10), and a similar regulatory mechanism (34). It would be interesting to test whether this promoter is heterogeneous and whether its activity impacts T6SS assembly. Another possibility is that T6SS assembly/activation is regulated by an accessory protein. Recent studies have identified accessory proteins that play a role in prey sensing and signalling to optimize T6SS assembly (56–58).

The T6SS is an important weapon used by bacteria to invade a niche. The observation of several gain and loss in certain species underlies a huge fitness cost for the T6SS at long term (53,59–61). The fine-tuned regulation of T6SS expression and activity thus appears to be key in the population development. However, constitutively expressed T6SS can be observed, notably in non-pathogenic strains, underlying the importance of considering the bacterium lifecycle to understand how is modulated the T6SS activity. Phenotypic heterogeneity of the T6SS in EAEC may represent an efficient strategy to enhance fitness in polymicrobial environment. *In vivo* approaches should determine whether this phenomenon is essential for pathogenesis. Since T6SS expression and activity is specific from species to species, further studies on its structure, activity and regulation in the diversity would bring information on its ecological function and on how bacteria adapt and shape communities.

## MATERIAL AND METHODS

### Bacterial strains and culture conditions

Bacteria and plasmids used in this study are listed in **Table S1**. *E. coli* strains were routinely grown in LB medium (iron-rich conditions) or in *sci1* inducing minimal medium (SIM, iron-starved conditions; M9 minimal medium supplemented with glycerol 0.2%, vitamin B1 200 μg/mL, casaminoacids 40 μg/mL, MgSO_4_ 1 mM and CaCl_2_ 0.1 mM). Media were supplemented with appropriate antibiotics (chloramphenicol 40 µg/mL, kanamycin 50 µg/mL, ampicillin 100 µg/mL, gentamycin 15 µg/mL) or isopropyl-*β*-D-galactopyrannoside (IPTG, 0.1 mM) when necessary. Cultures were grown at 37°C with agitation at 180 rpm. Chloro-Phenol-Red-Galactopyrannoside (CPRG, 2 mM) was prepared in distilled water.

### Plasmid construction

Plasmid and oligos used in this study are listed in **Table S2** and **Table S3**, respectively. PCR amplification was performed with a Biometra thermocycler using the Q5 polymerase (New England Biolabs). The pKO3-P*sci1* vector were obtained by restriction-ligation. Briefly, a DNA fragment containing the P*sci1* promoter (from - 500 pb to + 500 bp relative to the −10 element) was amplified from EAEC 17-2 genomic DNA with oligonucleotides carrying *Bam*HI and *Sal*I restriction sites at 5’ and 3’, respectively. Purified fragment and plasmid were digested by BamHI and SalI (New England Biolabs) for 3 h at 37°C and ligated with the T4 DNA ligase (New England Biolabs) for 1 h at room temperature. Point mutations of the promoter were engineered by quick-change mutagenesis. Briefly, pairs of complementary oligonucleotides containing the desired substitution were used to amplify the whole pKO3-P*sci1* plasmid. Amplicons were then treated with DpnI to eliminate template plasmids prior to transformation into DH5*α* competent cells. All pKO3-P*sci1* variants were verified by PCR and DNA sequencing (Eurofins Genomics).

### Strain construction

The fluorescence transcriptional reporter was constructed by *λ*-red-mediated recombination (62) using plasmid pKOBEG (63). Briefly, electrocompetent cells were transformed with pKOBEG and prepared for transformation with PCR products. The pKD4-Nt-sfGFP was used to amplify a fragment comprising the sfGFP gene and the kanamycin cassette flanked by 50 pb complementary to the upstream and downstream regions of the insertion site. PCR products were purified (PCR and DNA Clean up, Promega) and electroporated into pKOBEG-induced cells. Transformants clones were selected on kanamycin plates, checked by colony PCR before and sequencing of a PCR product encompassing the site of insertion to verify proper insertion at the desired position. The kanamycin cassette was then removed by using vector pCP20 (62). Directed mutagenesis was performed by the allelic exchange method (64) using the pKO3 vector carrying *sacB* which expression is toxic in the presence of sucrose. Briefly, electrocompetent cells were transformed with the pKO3-P*sci1* variants and plated at 30°C for plasmid replication. A few clones are re-streaked at 37°C for selecting clones with an integrated plasmid. The second event of recombination was obtained by re-streaking clones on LB agar plates supplemented with 5% of sucrose at 37°C. Clones were isolated on LB agar plate and randomly sent for sequencing to check the presence of the desired mutation.

### Competition assay

Competition assays using EAEC 17-2 and derivatives strains as a attacker and W3110 Kan^R^ as a recipient were performed as follow: strains were cultured overnight and diluted in a fresh SIM culture until optical density at *λ*=600 nm (OD_600_) of 0.8. Then, attacker and recipient were mixed in a 4:1 ratio for quantitative measurement using the lysis-associated β-galactosidase assay (LAGA) and 1:4 for qualitative measurement using the survivors growth kinetics (SGK) (33). Mixes were spotted in SIM agar plate and incubated at 37°C for 4 h. Briefly, the qualitative test is based on the chlorophenol-red β-D-galactopyranoside (CPRG) conversion to chlorophenol red (CPR) chromogeneous product due to cell lysis and β-galactosidase release. The spot colour being dependent of the killing efficiency, the intensity of the coloration is used as readout to compare each condition. For the quantitative measure, spots were resuspended in LB supplemented with 10 μg/mL of kanamycin prior to be 10-fold diluted and 2-fold diluted in a 96-wells microplate to select and measure W3110 surviving recipient growth using a TECAN microplate reader. Growth parameters were extracted using the R package growthcurver plugin (65). Since the time to resume growth depends on the initial cell concentration, the mid exponential time point (Tmid) was used as a readout to measure the recipient mortality and to compare each condition. Competition assay using *Pseudomonas aeruginosa* PAO1 *ΔretS* as a attacker and EAEC 17-2 pBBR5-Gm^R^ as a recipient were performed as follow: after an overnight culture in LB, PAO1 was diluted 25 fold in the same medium and cultured until an OD_600_ of 2. One mL of the culture was harvested and concentrated at an OD_600_ of 10 after centrifugation. EAEC 17-2 and derivatives strains were diluted in SIM from an overnight culture in LB supplemented with 30 μg/mL of Gentamycin and harvested at an OD_600 of_ 0.8. Heterogeneity ratios were artificially recreated by mixing F2 (ON) and G123 (OFF) strains in different ratios (0, 20, 40, 60, 80 and 100% of ON cells). Attacker-recipient mixes were then prepared at a 1:1 ratio, 10 µL were spotted on a SIM agar plates, in triplicate, and incubated at 37°C for 5 h. Bacterial spots were then treated for the quantitative measurement as described above.

### Fluorescence Microscopy

Cells were grown in liquid culture as mentioned above before to be sampled and concentrated about 10 times. 2 µL were then spotted onto an agarose pad 2% poured in a gene frame. The microscopy slide was closed by a coverslip prior to be observed under the microscope. Microscopy images were taken with a Nikon Eclipse Ti2 equipped with an ORCA flash 4.0 digital camera (Hamamatsu). Fluorescence of the sfGFP and mCherry was acquired at an excitation wavelength of 488 nm and 561 nm, respectively. Images were analysed through MicrobeJ after a segmentation step by MiSiC (66) to extract single-cell fluorescence.

Then, the fluorescence distribution was plotted using R package ggplot2 and geom_density function (67). The coefficient of variation was calculated as the ratio between the standard deviation and the mean fluorescence of the full data set. It was used as the readout for the heterogeneity level with the strain C-GFP-K. Since the T6SS sheath gives the brighter pixel in a cell compared to a diffuse fluorescence, the presence of a sheath was determined by the ratio between the max fluorescence and the mean fluorescence that was >2. The frequency of cells with sheaths (T6SS^+^) determined by this method was used as a proxy to measure the heterogeneity with the strain B-GFP.

### Flow cytometry and cell sorting

Influx v7 Cell Sorter (Becton, Dickinson and Company, Franklin Lakes, NJ, USA) was equipped with a stream-in-air 70 µm nozzle and a blue 488-nm Sapphire OPS laser (400 mW). The 488-nm laser was used for the analysis of the FSC (488/10 nm band pass filter, PMT1) which provides information related on cell size, the SSC (trigger signal, 488/10 nm band pass filter, PMT2) which provides information on cell density and the GFP fluorescence (530/40 nm band pass filter, PMT3). Light was detected with PMTs (Hamamatsu R3896 PMTs in C6270 sockets). As sheath FACSFlow buffer (Becton, Dickinson and Company) ran at 33 psi with an event rate of approx. 3,000 events s^-1^. The instrument was calibrated daily with 1 μm blue and 2 μm yellow-green FluoSphere beads (both Thermo Fisher Scientific, F-8815 and F-8827, Waltham, MA, USA) in the linear range. For calibration in the logarithmic range and for spiking into the sample, 0.5 μm and 1 µm yellow-green beads (both Thermo Fisher Scientific, F-8813 and F-13081, Waltham, MA, USA) were used. Measurement was stopped after 50,000 events were recorded in the FSC against SSC cell gate. Fluorescence-activated cell sorting (FACS) was performed as in (68) with the most accurate sorting mode ‘1.0 drop pure’ and at a maximum sorting speed of 3,000 cells s^-1^. 2×10^6^ cells per gate were sorted for microscopy and stored with 15 % glycerol at −80 °C until further experiments. 2D-plots were created using FlowJo V10 (Becton, Dickinson and Company). In addition, proportions of cells were analyzed in 2D plots FSC vs green fluorescence in the GFP cell gate equal to 100% with respect to the 3 gates assembler, producer (ON) and non-producer (OFF) for the 4 strains EAEC 17-2, B-GFP, C-GFP-K and C-GFP-K F2, respectively (**Figure S4**). For the calculation, the beads in the GFP cell gate were subtracted.

### Single-cell experiments

Single-cell experiments were performed using the Ibidi sticky-Slide VI 0.4 device as follow: BD agar 2% was poured into the channel before to be dug by cutting the agar to build a new channel for fluid passage; the cell suspension was spotted on the agar and covered with a slide; the input well was filled with 80 µL of culture medium and the device was incubated at 37°C in the hand-made thermostable chamber under the microscope. After 4 h of incubation, images of growing microcolonies are taken with settings described above.

### Wax moth infection assays

*Galleria mellonella* was used as an animal model of infection. Prior to infection, larvae underwent nutrient starvation for 48 h at 28°C in the dark. EAEC 17-2 F2 and G123 were used to mimic a homogeneous ON or OFF population, respectively, and compared to the WT. Strains were grown in LB until OD_600_ 0.8. Then, starved larvae were infected by natural ingestion of 2×10^5^ CFU/mL and were incubated for 24 h at 28°C in the dark. After 24 h, each larva was washed for 10 sec in sterile water and sacrificed to collect intern organs. The content was resuspended in 500 µL of PBS with 6 glass beads and destroyed using the FastPrep-24 5G machine (MP biomedicals) during 30 sec at power 4.5. 100 µL of lysate were serially diluted and plated on agar plate supplemented with 20 µg/mL of kanamycin for EAEC recovery. The experiment was performed in triplicate.

## ACKNOWLEDGEMENTS

We thank Julien Brillard and Alain Givaudan (DGIMI, INRAE Montpellier, France) for help regarding *Galleria mellonella* infection assays, Marie Grandjean and Corine Sebban-Kreuzer for the PAO1*ΔretS* strain and discussions, members of the Cascales laboratory for discussions, and Moly Ba, Isabelle Bringer, and Annick Brun for technical assistance. This work was supported by the CNRS, the Aix-Marseille Université and by grants from the Agence Nationale de la Recherche (ANR-20-CE11-0017), the Fondation pour la Recherche Médicale (DEQ20180339165), the Fondation Bettencourt-Schueller to EC and the German Research Association (503905203) to SM.

## Data availability

All data are shown in the manuscript or in the supplemental material.

## Declaration of competing interest

The authors declare that they have no known competing financial interests or personal relationships that could have influenced the work reported in this paper.

## Notes

### Competing Interest Statement

The authors have declared no competing interest.

