## Supplemental Information for "Type VI secretion phenotypic heterogeneity ensures trade-off between antibacterial activity and resistance in Enteroaggregative *E. coli*"

### Table S1. Strains used in this study

| Name | Strain | Description | Source |
|---|---|---|---|
| WT | Entero-aggregative *E. coli* 17-2 | Wild-type strain | Lab collection |
| *sci1* | 17-2 Δ*sci1* | Deletion of the T6SS *sci1* gene cluster | Lab collection |
| *lacZ* | 17-2 *lacZ*::kan | 17-2 with $Kan^R$ cassette inserted at *lacZ* locus | This study |
| *sci1 lacZ* | 17-2 Δ*sci1 lacZ*::kan | 17-2 Δ*sci1* with $Kan^R$ cassette inserted at *lacZ* locus | This study |
| B-GFP | 17-2 TssB-GFP | sfGFP fused to *tssB* | Zoued *et al.,*2013 |
| C-GFP-K | 17-2 P*sci1*-GFP | sfGFP inserted between *tssC* and *tssK* genes | This study |
| G1 | 17-2 GATC1* | P*sci1* GATC1->GGTC | This study |
| G2 | 17-2 GATC2* | P*sci1* GATC2->GGTC | This study |
| G3 | 17-2 GATC3* | P*sci1* GATC3->GGTC | This study |
| G1G2 | 17-2 GATC1* GATC2* | GATC1 and 2 double mutant | This study |
| G123 | 17-2 GATC1* GATC2* GATC3* | GATC triple mutant | This study |
| F1 | 17-2 Fur box1* | P*sci1* Fur box1 mutant | This study |
| F2 | 17-2 Fur box2* | P*sci1* Fur box2 mutant | This study |
| F1F2 | 17-2 Fur boxes* | Fur box double mutant | This study |
| PAO1Δ*retS* | *P. aeruginosa* | H1-T6SS upregulation | C. Kreuzer |
| DH5a | *E. coli* K-12 DH5a | Wild-type *E. coli* K12, used for cloning | NEB |
| W3110 | *E. coli* K-12 W3110 | Wild-type *E. coli* K12, used as recipient strain | K. Postle |

### Table S2. Plasmids used in this study

| Plasmid | Composition | Description |
|---|---|---|
| pJET | $Amp^R$ | Cloning vector for quick change mutagenesis |
| pKO3 | SacB, $Cm^R$, *ori*$^{ts}$ | Suicide vector for chromosomal site-directed mutagenesis |
| pKO3-P*sci1* | P*sci1* DNA fragment cloned into pKO3 | Suicide vector with DNA fragment encompassing P*sci1* promoter |
| pKO3-P*sci1*-F1 | P*sci1* DNA fragment with F1 mutation cloned into pKO3 | pKO3-P*sci1* with F1 mutation |
| pKO3-P*sci1*-F2 | P*sci1* DNA fragment with F2 mutation cloned into pKO3 | pKO3-P*sci1* with F2 mutation |
| pKO3-P*sci1*-G1 | P*sci1* DNA fragment with G1 mutation cloned into pKO3 | pKO3-P*sci1* with G1 mutation |
| pKO3-P*sci1*-G2 | P*sci1* DNA fragment with G2 mutation cloned into pKO3 | pKO3-P*sci1* with G2 mutation |
| pKO3-P*sci1*-G3 | P*sci1* DNA fragment with G3 mutation cloned into pKO3 | pKO3-P*sci1* with G3 mutation |
| pKO3-P*sci1*-G123 | P*sci1* DNA fragment with G1, G2 and G3 mutations cloned into pKO3 | pKO3-P*sci1* with G1, G2 and G3 mutations |
| pKD4 | FRT-$Kan^R$-FRT | Template for lambda-red recombination |
| pKOBEG | Rec genes, $Cm^R$, *ori*$^{ts}$ | Plasmid-encoding lambda-red system |
| pBAD33 | Plac, CmR | Expression vector |
| pPROBE-gfp-AAV | gfp-AAV, $Kan^R$ | Transcriptional reporter plasmid with unstable GFP |
| pPROBE-gfp-AAV-Fur | P*fur*, gfp-AAV, $Kan^R$ | Unstable GFP under the control of P*fur* promoter |
| pPROBE-gfp-AAV-Sci1 | Ps*ci1*, gfp-AAV, $Kan^R$ | Unstable GFP under the control of Ps*ci1* promoter |
| pBBR-MCS5 | $Gm^R$ | Broad Host Range plasmid with gentamycin gene |

### Table S3. Oligonucleotides used in this study

| | |
|---|---|
| **Cloning into pJET/pKO3** | |
| 5-pKO3-Psci-1 | TATCGGATCCGCCGGCAGGGATTTCG |
| 3-pKO3-Psci-1 | ACGGTCGACCGGCATGCGCTTCAG |
| **Quick-change mutagenesis** | |
| 5-G1 | CAGATCTTCGTCCTATAATGGTCAAAATTAAATCAGTGCACAAG |
| 3-G1 | CTTGTGCACTGATTTAATTTTGACCATTATAGGACGAAGATCTG |
| 5-G2 | CTGTTATATTGAGATTTTTCAGGTCTTCGTCCTATAATG |
| 3-G2 | CATTATAGGACGAAGACCTGAAAAATCTCAATATAACAG |
| 5-G1-2 | CTGTTATATTGAGATTTTTCAGGTCTTCGTCCTATAATGGTCAAAATTA AATCAGTGCACAAGGG |
| 3-G1-2 | CCCTTGTGCACTGATTTAATTTTGACCATTATAGGACGAAGACCTGAA AAATCTCAATATAACAG |
| 5-G3 | CTGATTATTTGCATTATATCGGTCGATGTATCTGTTATATTG |
| 3-G3 | CAATATAACAGATACATCGACCGATATAATGCAAATAATCAG |
| 5-F1 | CGTCCTATAATGATCAAAAGGGAATCAGTGCACAAGGGGAGGC |
| 3-F1 | GCCTCCCCTTGTGCACTGATTCCCTTTTGATCATTATAGGACG |
| 5-F2 | TAAAGTCCTGATTATTTGCAGGGTATCGATCGATGTATCTGTT |
| 3-F2 | AACAGATACATCGATCGATACCCTGCAAATAATCAGGACTTTA |
| **Cloning into pPROBE** | |
| 5-pPROBE-Psci-1-SalI | TATCGTCGACTCATTGCATTTTGTTGGGCCCT |
| 3-pPROBE-Psci-1-EcoRI | ACGGAATTCGGCTCTCTCCTGTGAAACCTGC |
| 5-pPROBE-P*fur*-EcoRI | TATCGTCGACTCTCGGTCTGGCTATCGACG |
| 3-pPROBE-P*fur*-SalI | ACGGAATTCGCGGAATCTGTCCTG |
| pKD4-Nt-sfGFP | |
| 5-tssC-sfGFP-tssK | TGGACGTCAGCCTGTCACTGGTTTCGCAGATGCCGAAGGCAAAAGCG TAACGATTGTGTAGGCTGGAGCTGCTTCGAAGTTCCTATAC |
| 3-tssC-sfGFP-tssK | GCCCCGTCTTCCCATAATGGGCGATAAATCTTCATTTCCCGACACCTG CCTTACCCTCCGCCGGCCGCTGC |

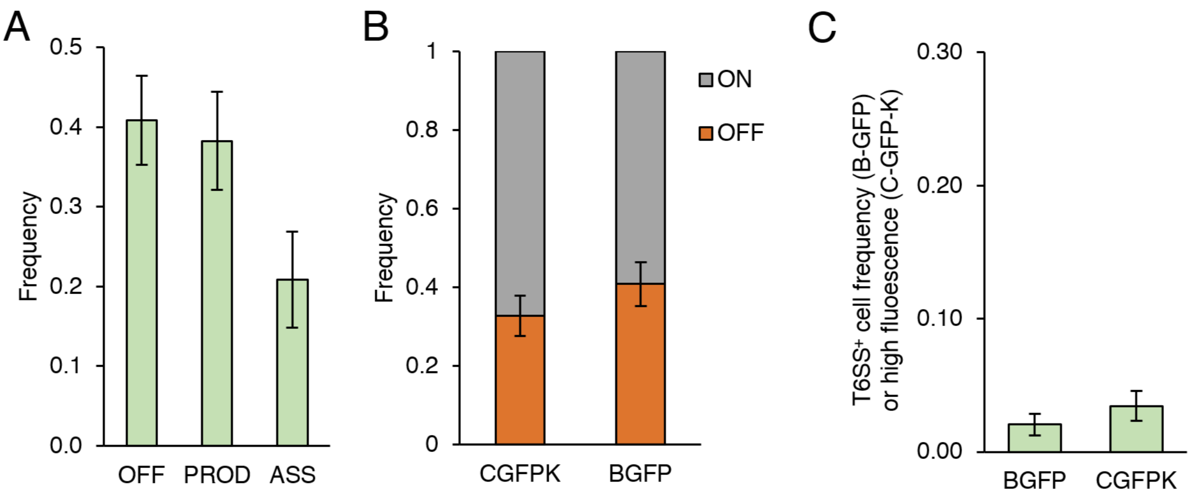


**Figure S1. Phenotypic heterogeneity of T6SS expression and assembly. A**. Frequency of cell with no sheath (OFF), TssB production (PROD) or assembled T6SS (ASS) in a clonal population of EAEC B-GFP. 5,450 cells were analysed, from 3 independent replicates. **B**. Frequency of ON and OFF cells in a clonal population of C-GFP-K (11,760 cells analysed) or B-GFP (5,450 cells analysed). Data shown are from 3 independent replicates. **C**. T6SS$^{+}$ cell frequency (B-GFP) or high fluorescence (C-GFP-K) in iron-rich LB medium. 1,518 B-GFP cells and 1,945 C-GFP-K cells were analysed from 3 independent replicates.

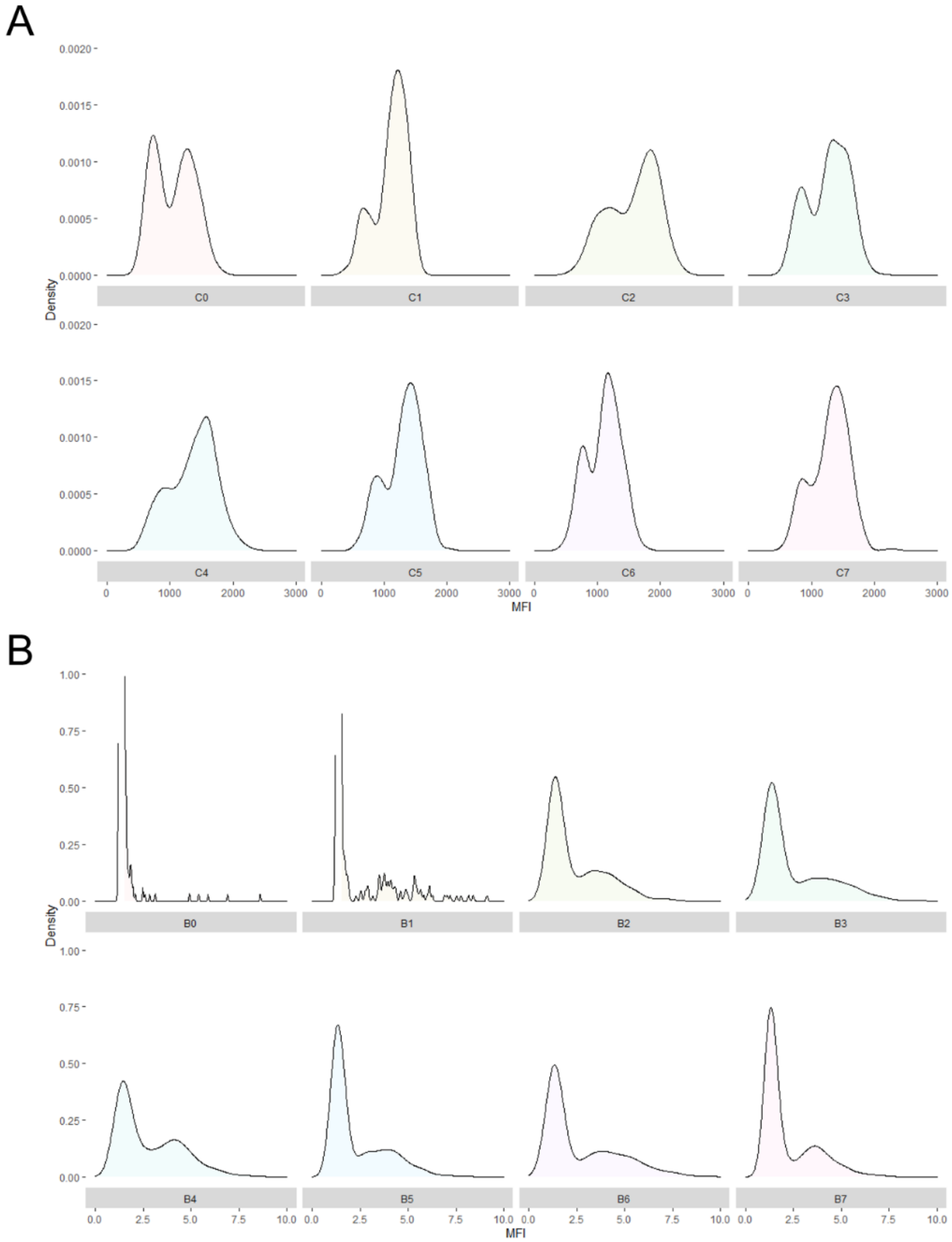


**Figure S2. Phenotypic heterogeneity of T6SS expression and assembly over generations. A.** Representative distribution (Density) of the mean fluorescence intensity (MFI) of C-GFP-K cells in the populations over seven generations (C0 to C7) (*n*>400 cells for each generation). **B.** Representative distribution of max/mean fluorescence intensity ratio (sheath detection) of B-GFP cells in the population over seven generations (B0 to B7) (*n*>300 cells for each generation).

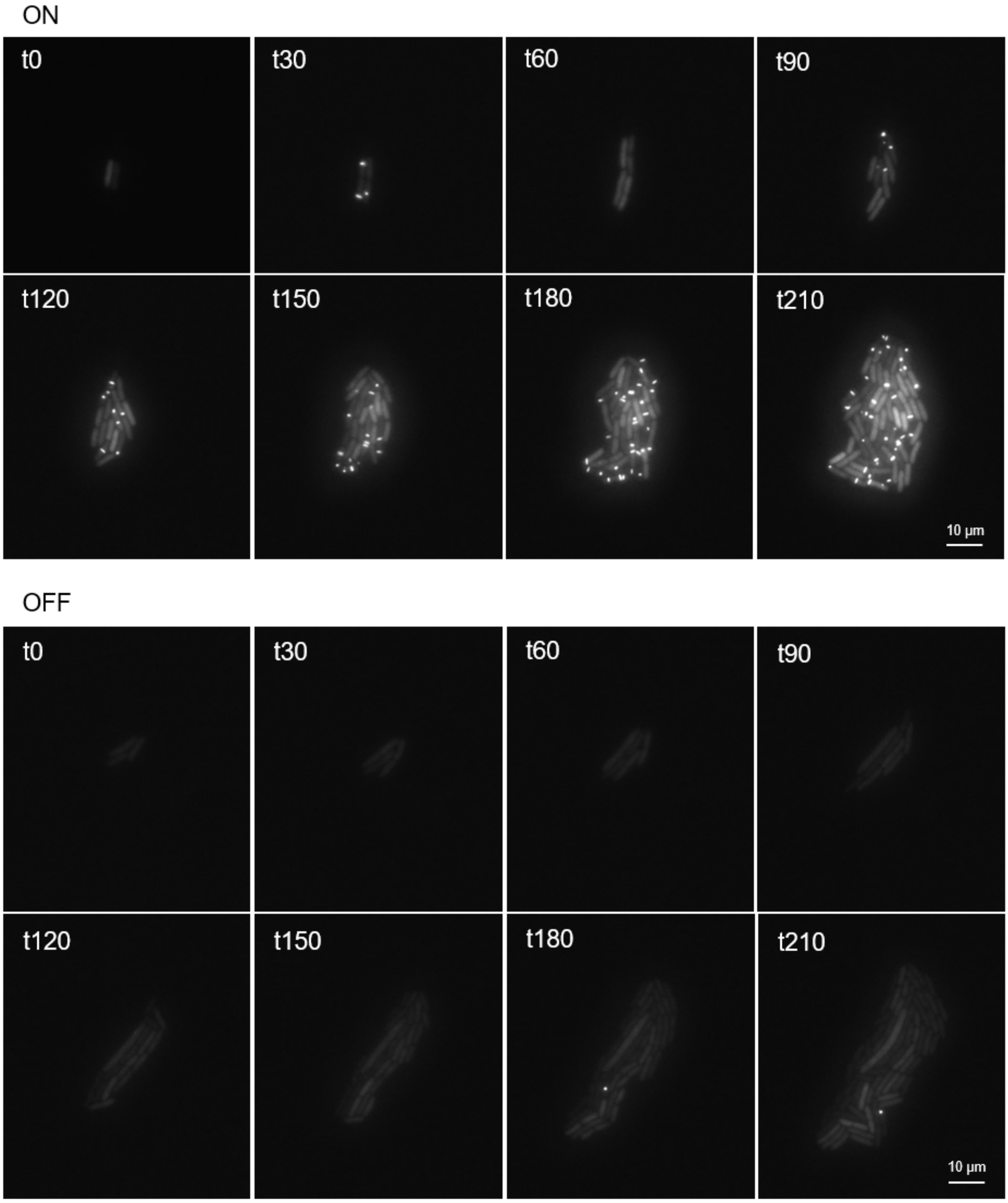


**Figure S3. Representative time-lapse recordings of single-cell experiment from a clonal B-GFP cell.** Single-cell imaging of a ON (top) and OFF (bottom) cell grown in a microfluidic device fed with SIM medium. Images were taken every 30 minutes from t0.

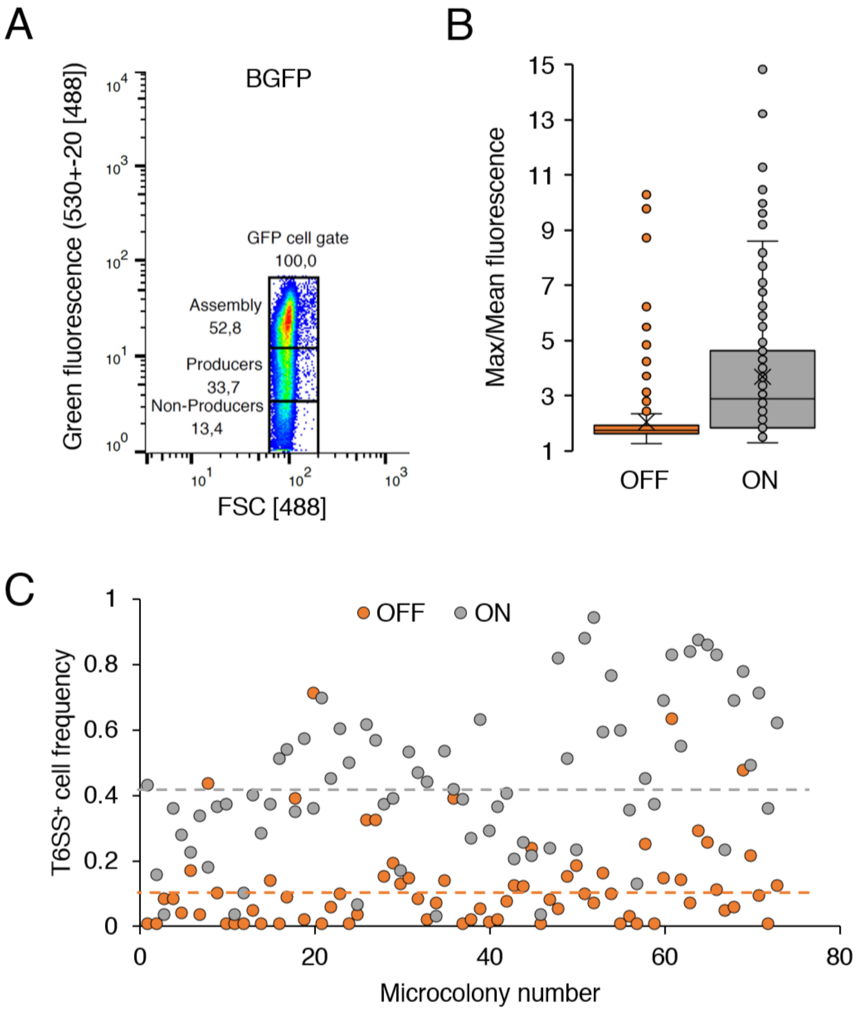


**Figure S4. Cell sorting of subpopulations. A**. Distribution of the mean fluorescence of a TssB-GFP population determined by flow cytometry at 6h of culture after 1/100e dilution of a preculture in SIM+LB 10%. Non producers gate was defined with the autofluorescence of EAEC 17-2 strain; Assembly gate was defined with the fluorescence distribution of C-GFP-K F2; Producers gate was defined as the aera of medium intensity between Non-producers and Assembly gates. **B**. Quality control of the cell sorting of ON (assembler and producer) and OFF (non-producer) subpopulations. The presence of a sheath is determined by a max and mean fluorescence ratio >2. Each point is a cell (*n*=175 and 184 OFF and ON cells analysed from 2 independent samples). **C**. T6SS$^{+}$ cell frequency in microlonies from single-cell experiments of sorted ON and OFF subpopulations. Each dot is a microcolony. The overall average frequency is shown as a dotted line with the corresponding colour. More than 3,400 cells were analysed from 73 microcolonies from 6 independent replicates.

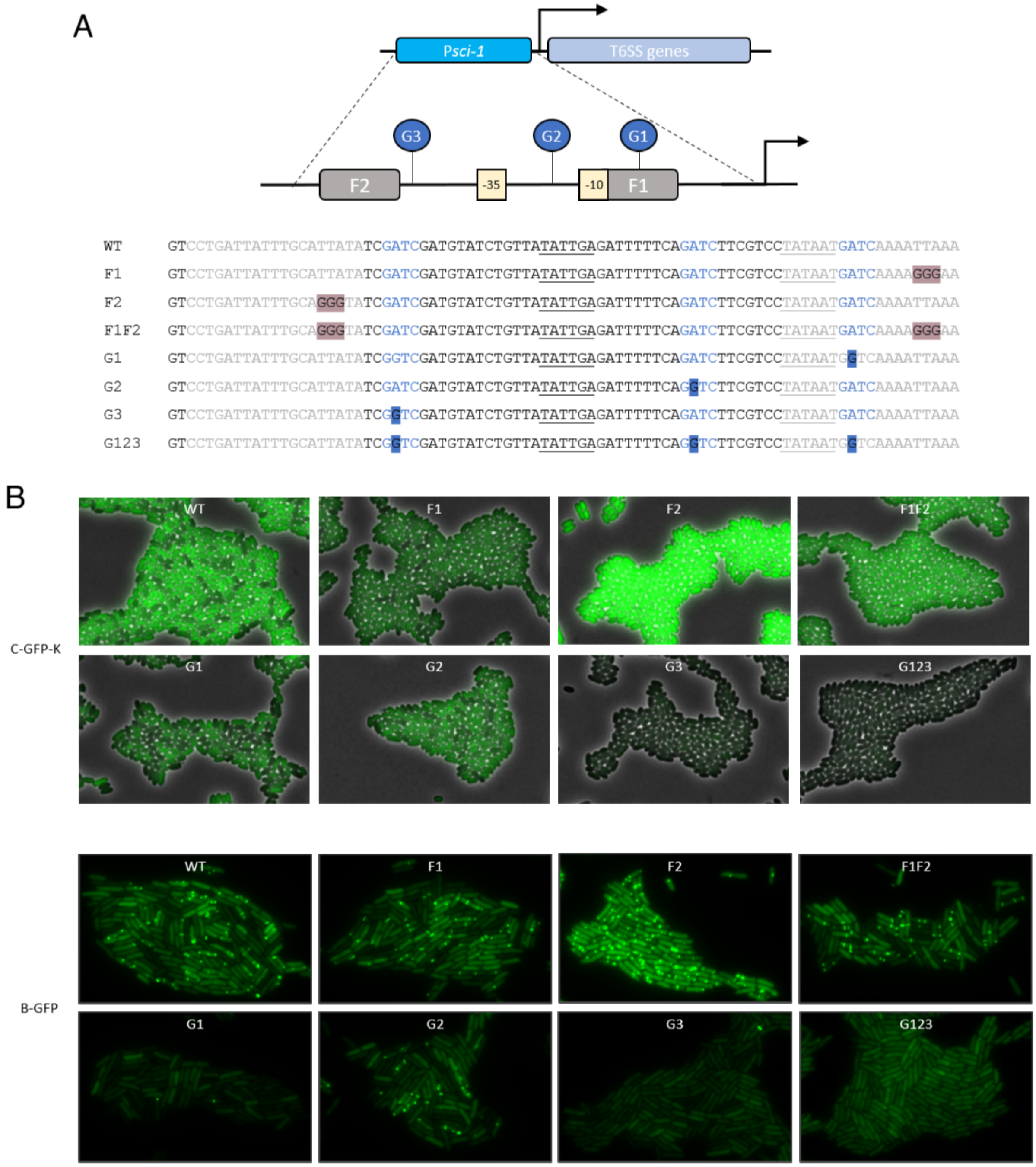


**Figure S5. Impact of promoter point mutations on T6SS$^{+}$ cell frequency (B-GFP) and T6SS expression (C-GFP-K). A**. Sequence alignment of WT and variant P*sci1*. Fur boxes sequences are coloured in grey, GATC sites in blue and -10 and -35 elements are underlined. Fur boxes were mutated by the substitution of 3 bases (GGG highlighted in grey) to break the palindrome. GATC sites were mutated by the substitution of the Adenine by a Guanosine (G highlighted in blue) to prevent methylation. **B**. Representative fields of the phenotype caused by mutations. C-GFP-K images are merged channels of phase and fluorescence set up with the same minimal and maximal signal scale. B-GFP images are only fluorescence images to better observe T6SS apparatus. G3 and G123 are overexposed to distinguish cells.

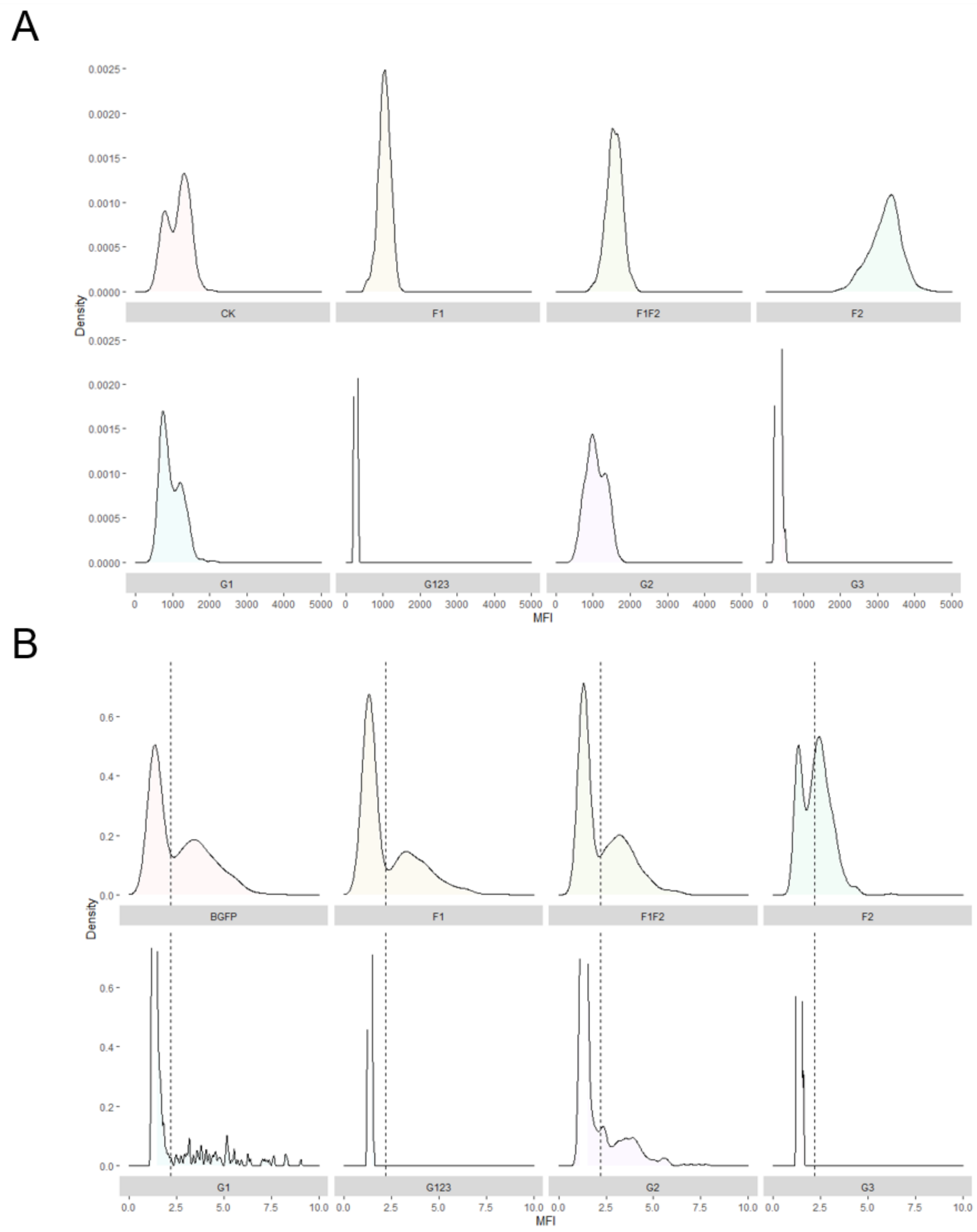


**Figure S6. Phenotypic heterogeneity of the T6SS expression and assembly of promoter variants.** **A**. Representative distribution (Density) of the mean fluorescence (MFI) of C-GFP-K populations in promoter variants. >500 cells per strains were analysed, from 3 independent replicates. **B**. Representative distribution of max/mean fluorescence ratio (sheath detection) of TssB-GFP populations in promoter variants. >500 cells per strains were analysed, from 3 independent replicates.

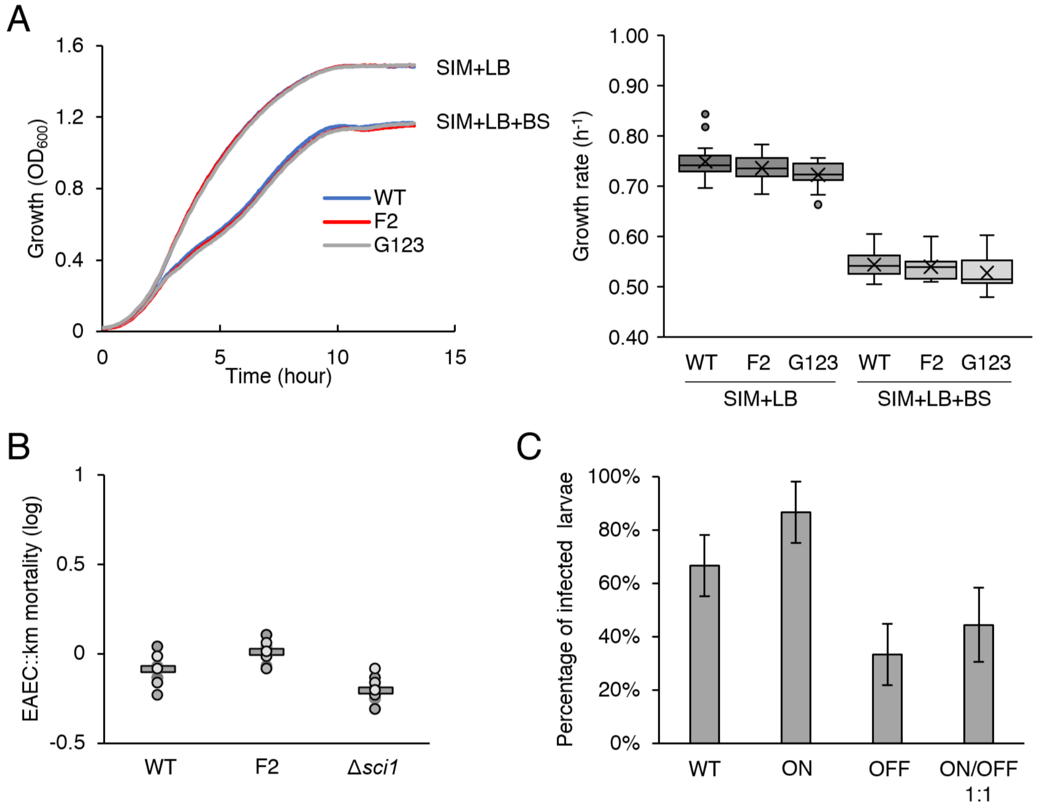


**Figure S7. Potential roles of T6SS heterogeneity. A**. Growth curve (left) and growth rate (right) of EAEC WT, F2 and G123 derivatives in SIM+LB 10% or SIM complemented with bile salts (SIM+LB+BS). The growth rate was calculated using growthcurver package in R studio (65). Data represent 3 independent replicates. **B**. Competition assay between EAEC WT, F2 or Δ*sci1* with EAEC carrying kanamycin resistance (EAEC::km). Mortality was measured by the survivors kinetic growths method on 2 independent replicates. **C**. Infective success of EAEC WT, ON homogeneous F2 strain, OFF Δ*sci1* mutant strain or a 1:1 mix of F2 and Δ*sci1* in the *Galleria mellonella* host. Infective success is determined as a recovery of detectable CFU on agar plates. Data are the mean of 3 independent replicates, each with 15 larvae.

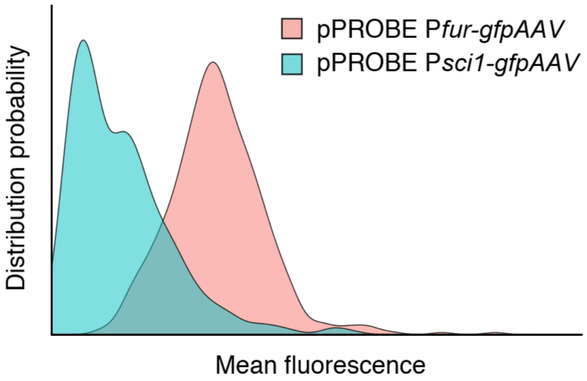


**Figure S8. Representative distribution profile of P*fur* and P*sci1* activity in EAEC carrying pPROBE-*gfpAAV* fused with the corresponding promoter grown in SIM+LB 10%.** The graph depicts a heterogeneous distribution of P*sci1* activity but a homogeneous P*fur* activity in the populations grown in the same conditions.
